# Timing of dense granule biogenesis in asexual malaria parasites

**DOI:** 10.1101/2023.06.19.545557

**Authors:** Tansy Vallintine, Christiaan van Ooij

## Abstract

Malaria is an important infectious disease that continues to claim hundreds of thousands of lives annually. The disease is caused by infection of host erythrocytes by apicomplexan parasites of the genus *Plasmodium*. The parasite contains three different apical organelles – micronemes, rhoptries and dense granules – whose contents are secreted to mediate binding to and invasion of the host cell and the extensive remodelling of the host cell that occurs following invasion. Whereas the roles of micronemes and rhoptries in binding and invasion of the host erythrocyte have been studied in detail, the role of dense granules (DGs) in *Plasmodium* parasites are poorly understood. They have been proposed to control host cell remodelling through regulated protein secretion after invasion, but many basic aspects of the biology of DGs remain unknown. Here we describe DG biogenesis timing for the first time, using RESA localisation as a proxy for DG formation timing. We show that DG formation commences approximately 37 minutes prior to schizont egress, as measured by the recruitment of the DG marker RESA. Furthermore, using a bioinformatics approach, we aimed to predict additional cargo of the DGs and identified the J-dot protein HSP40 as a DG protein, further supporting the very early role of these organelles in the interaction of the parasite with the host cell.

## INTRODUCTION

Malaria is caused by invasion, remodelling and lysis of host red blood cells by parasites of the genus *Plasmodium*. Invasion of the host cell is controlled by the regulated secretion of proteins from three specialised secretory organelles: micronemes, rhoptries and dense granules (DGs). The presence of three distinct apical organelles allows for compartmentalisation of proteins with specific functions and temporal regulation of protein discharge for rapid and efficient invasion and modification of the host cell (Cao et al., 2009; Carruthers & Sibley, 1997; Gubbels & Duraisingh, 2012; Kats et al., 2008; Singh et al., 2010). Micronemes are the first to secrete their contents (Singh et al., 2010) and primarily control host cell recognition, attachment, and invasion through release of proteins such as erythrocyte binding antigen 175 (EBA-175) (Adams et al., 1992; Camus & Hadley, 1985; Kim et al., 1992) and apical membrane antigen-1 (AMA-1) (Bannister et al., 2003; Mitchell et al., 2004; Peterson et al., 1989; Triglia et al., 2000). The rhoptries are second to discharge their contents, which includes lipid whorls in addition to proteins (Singh et al., 2010; Stewart et al., 1986); based on the timing of discharge it was thought that a subset of rhoptry proteins and the lipids secreted from the rhoptries initiate and support parasitophorous vacuole (PV) formation and invasion (Bannister & Mitchell, 1989; Cao et al., 2009; Ladda et al., 2001). This has been supported by the finding that the rhoptry-associated protein (RAP) complex facilitates parasite growth and survival within the host cell after invasion, with conditional knockdown of RAP components resulting in structural deformity of the PV membrane (PVM), delayed intraerythrocytic development and decreased parasitaemia in mouse models (Ghosh et al., 2017). Although the content of apical organelles appears to be segregated along functional lines, studies in *Plasmodium* parasites and *Toxoplasma* parasites have revealed that cooperation between microneme and rhoptry proteins occurs to allow parasite invasion of the host erythrocyte (Alexander et al., 2005; Besteiro et al., 2009). Binding of the microneme protein AMA1 and the rhoptry neck protein RON2 triggers formation of the moving junction complex (MJ) (Cao et al., 2009; Lamarque et al., 2011). The MJ functions as an interface between the membranes of the invading parasite and the host cell through which the parasite passes into the host cell during invasion. Blockage of the AMA1 binding site of RON2 inhibits formation of the MJ and parasitophorous vacuole (Cao et al., 2009; Srinivasan et al., 2011).

DGs are the last organelle to secrete their contents (Culvenor et al., 1991; Riglar et al., 2011; Torii et al., 1989) and are speculated to be required for the remodelling of the host cell after invasion, thereby allowing parasite survival and replication through the asexual stages (Carruthers & Sibley, 1997; De Koning-Ward et al., 2016). However, little is known about DGs in *Plasmodium* parasites; only few DG proteins have been identified in *Plasmodium falciparum*, whilst over 40 have been described in the related apicomplexan parasite *Toxoplasma gondii* (Griffith et al., 2022; Ming et al., 2018), in which DGs have been studied more extensively than in *Plasmodium* parasites. The few known *P. falciparum* DG proteins enable transport of parasite effector proteins into the erythrocyte (Elsworth et al., 2014; Garten et al., 2018; Ho et al., 2018) or are exported and subsequently mediate alterations of the biochemical and biophysical properties of the host cell (Pei et al., 2007). The only known contents of *Plasmodium* DGs are the five core proteins of the PTEX (*Plasmodium* translocon of exported proteins) – EXP2, HSP101, PTEX150, PTEX88 and TRX2 – an essential protein complex that transports parasite effector proteins across the PVM into the host cell (Beck et al., 2014; Bullen et al., 2012; Charnaud et al., 2018; De Koning-Ward et al., 2009; Elsworth et al., 2014); PV1, which interacts with exported proteins and the PTEX (Chu et al., 2011; Morita et al., 2018); EXP1, a transmembrane protein in the parasitophorous vacuole membrane (Gunther et al., 1991; Iriko et al., 2018; Simmons et al., 1987) that has an important role in nutrient uptake across the PVM through its effect on the localisation of EXP2 (Garten et al., 2018; Gunther et al., 1991; Ho et al., 2018; Mesén-Ramírez et al., 2019); P113, a GPI-linked protein that associates with the PTEX and with PVM and exported proteins (Bullen et al., 2022; Miyazaki et al., 2021); LSA3, the function of which is unknown (Morita et al., 2017); and RESA (ring-infected erythrocyte surface antigen), which binds to and stabilises spectrin tetramers below the erythrocyte surface, thereby reducing host cell deformability (Culvenor et al., 1991; E. Da Silva et al., 1994; Diez-Silva et al., 2012; Pei et al., 2007; Diez Silva et al., 2005) (Table 1). The recently published Dense Granule Protein Database (DGPD) (http://dgpd.tlds.cc/DGPD/index/), which provides information on identified and predicted DG proteins of Apicomplexan parasites, lists an additional 4 PHIST proteins and a ClpB1 orthologue as putative DG proteins (Hu et al., 2022), although the ClpB1 orthologue has also been identified previously as an apicoplast protein (El Bakkouri et al., 2010) (Table 1). Hence, based on the function of the known DG proteins, this organelle appears to facilitate the modification of the host cell into a hospitable environment for parasite growth. For example, the increased cell rigidity caused by RESA allows infected cells to avoid filtration by the spleen by sequestering cells in the microvasculature whilst also increasing heat shock resistance, allowing parasites to survive the increased temperatures of febrile episodes and inhibit further invasion by other merozoites (E. Da Silva et al., 1994; Diez-Silva et al., 2012; M. D. Silva et al., 2005). The timing of DG protein secretion immediately following invasion, the extensive nature of erythrocyte remodelling at that time and the greater number of DG proteins found in other Apicomplexan parasites all suggest that DGs of *Plasmodium* parasites likely contain many more exported proteins than those currently identified (Rezaei et al., 2019). This theory is further supported by the late-stage transcriptional profiles of 90 genes encoding putative exported proteins containing a *Plasmodium* Export Element (PEXEL) that are predicted to be exported into the host erythrocyte (Marti et al., 2004). This demonstrates that in *Plasmodium* parasites, exported proteins are synthesised very late in the life cycle (Hallée et al., 2018; Tomavo, 2014).

**Table 1.**
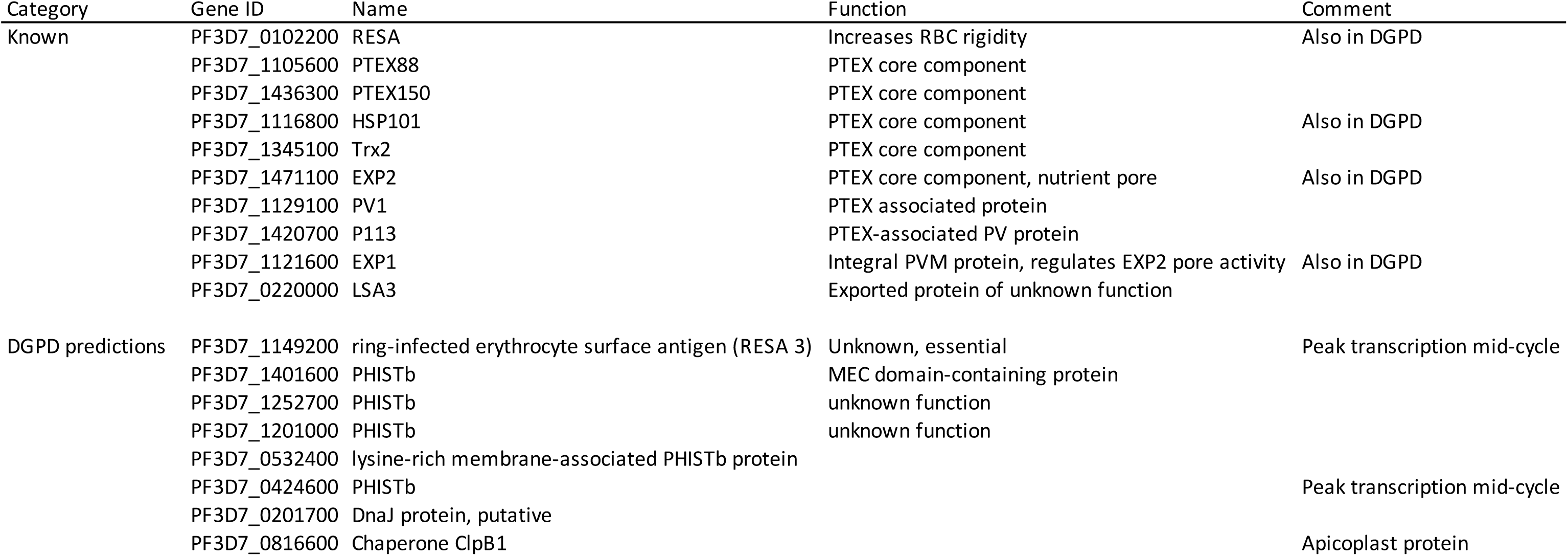
Known and predicted *Plasmodium falciparum* dense granule proteins

Despite the important role of DGs, many basic aspects of DG formation in *Plasmodium* parasites remain unknown, hampering the study of this important organelle. In this study we aimed to determine the timing of DG biogenesis and identify previously undescribed DG proteins. In order to investigate DG biogenesis timing, we generated parasites expressing RESA fused to mNeonGreen, allowing DG biogenesis to be observed by live video microscopy. Using this knowledge about DG biogenesis, we applied a bioinformatics approach to identify additional DG proteins.

## MATERIALS AND METHODS

### Parasite culture

*P. falciparum* erythrocytic stage parasites of strain 3D7 were cultured in human erythrocytes (UK National Blood Transfusion Service and Cambridge Bioscience, UK) at 3% haematocrit and 37°C, 5% CO_2_ in RPMI-1640 medium (Life Technologies) supplemented with 2.3 g/L sodium bicarbonate, 4 g/L dextrose, 5.957 g/L HEPES, 50µM hypoxanthine, 0.5% AlbuMax type II (Gibco), and 2 mM L-glutamine (complete RPMI (cRPMI) according to established procedures (Crabb et al., 1999)).

### Transfection

Parasites were transfected using the schizont method (Collins et al., 2013). For each transfection, 20 µg of plasmid in 100 µl buffer was precipitated using 10 µl sodium acetate and 250 µl 100% ethanol overnight at −80°C. Precipitated DNA was washed with 70% ethanol and air dried in a sterile environment before re-suspension in 10 µl sterile TE buffer. The DNA for the transfections was resuspended in 10 µl TE and 100 µl AMAXA P3 buffer solution was added.

To prepare the parasites for transfection, two T-75 flasks of 3D7 parasite cultures containing predominantly late-stage schizonts were synchronised tightly using two rounds of density gradient centrifugation using 70% Percoll (GE Healthscience) and incubated at 37°C with 25 nM of the parasite egress inhibitor ML10 for 1.5 hours (Ressurreiçao et al., 2020) to maximise numbers of late-stage segmented schizonts. The parasites were then washed with cRPMI to remove ML10 and resuspended in 5 ml cRPMI, which was distributed over 5 microcentrifuge tubes (1 tube per transfection) and incubated at 37°C for 18 minutes. The cells were pelleted, re-suspended in the DNA/AMAXA buffer solution and subsequently transferred to AMAXA transfection cuvettes and transfected in the AMAXA electroporation device using the P3 Primary Cell transfection reagent programme. The electroporated cell suspension was transferred to T-25 cell culture flasks containing 300 µl packed RBC in 2 ml cRPMI and the released merozoites were allowed to invade in a shaking incubator. After 30 minutes 8 ml medium was added to each flask and the cultures were transferred to a non-shaking incubator. After 24 hours WR99210 was added to 2.5 nM to the parasites to select for transfectants. Transfected parasites were generally recovered after approximately 3 weeks. Integrants were selected with treatment with G418 at 600 µg/mL.

### Plasmids

We aimed to produce a plasmid encoding a fusion of mNeonGreen (mNG) to the 3’ region of RESA (PF3D7_0102200), followed by sequence encoding the T2A skip peptide and a neomycin resistance marker, toallows for selection of parasites with integrated plasmid using G418 (Birnbaum et al., 2017b, 2017a). First, the gene encoding GFP was removed from the plasmid pRESA-GFP (de Azevedo et al., 2012) by digestion with PstI and MluI and replaced with a 705 bp insert encoding mNG that was amplified using primers CVO550 and CVO551 (Supplementary Table 2) and digested with PstI and MluI for insertion into the pRESA-GFP backbone, producing plasmid pTV001. The genes encoding the T2A peptide and G418 resistance marker were added by amplification of a fusion of these two genes from JT02-01-31 (a kind gift of James Thomas, LSHTM) using primers CVO576 and CVO577 (Supplementary Table 2) and inserted at the 3’ end of the mNG fragment by digestion of pTV001 and the PCR fragment with MluI. The fragments were joined using In-Fusion (TaKaRa), producing plasmid pTV002, encoding a fusion of the 3’ 821 bp of the gene encoding RESA, the T2A peptide and G418 the resistance marker.

### Genomic DNA isolation

Erythrocytes were pelleted by centrifugation at 2,000 xg for 5 minutes and the resulting pellet was resuspended in an equal volume of 0.15% saponin in PBS to release the parasites from the erythrocytes. Parasites were pelleted by centrifugation at 13,000 xg for five minutes and the parasite DNA was isolated using the Monarch^®^ Genomic DNA Purification Kit (New England Biolabs).

### Immunoblotting

Late-stage segmented schizonts were isolated from highly synchronised cultures by flotation on a 70% Percoll gradient (Rivadeneira et al., 1983) and blocked with 25 nM ML10 until Giemsa smears showed only late-stage segmented schizonts. The schizonts were pelleted by centrifugation at 2,000 xg for five minutes, pellets were resuspended in SDS-PAGE gel-loading buffer and then heated to 98°C for 20 minutes.

Proteins were separated by SDS-PAGE on 12.5% gels at 200 V for 45 minutes and subsequently transferred to a nitrocellulose membrane at 0.1 A and 25 V for 50 minutes using a Bio Rad Transblot Turbo transfer system. The membranes were blocked with 5% milk powder dissolved in PBS-Tween for one hour at room temperature before incubation with either anti-mNG primary antibody diluted 1:1000 in PBS-Tween or HRP-linked anti-aldolase antibody (Abcam) diluted 1:5000 in PBS-Tween for one hour. After extensive washing, the blot probed with the anti-mNG antibody (ChromoTek) was incubated with HRP-linked secondary antibody (Bio Rad) for one hour at room temperature and after extensive washing with PBS both blots were developed with Clarity ECL Western blotting substrate (Bio Rad). Both were imaged on a Bio Rad ChemiDoc and further cropped and sized using Abobe Photoshop. Figures were produced using Adobe Illustrator.

### Video microscopy

Highly synchronous late-stage schizonts were isolated on a 70% Percoll gradient. Serial dilutions of schizonts in cRPMI containing 100 µM resveratrol were transferred to Ibidi poly-L-lysine µ-Slide VI^0.4^ channel slides. Slide wells were then sealed with petroleum jelly to prevent sample dehydration. Parasites were maintained at 37°C during transport and transferred to a pre-heated Okolab Microscope Incubator Cage with Gas Micro-Environmental Chamber and Air Heater for live imaging. Imaging was carried out at 37°C and in an atmosphere containing 5% CO_2_ on a Nikon Eclipse TE fluorescence microscope equipped with a Hamamatsu ORCA-flash 4.0 digital camera C11440 controlled using Nikon NIS-Elements version 5.3 software. Schizonts were imaged every five minutes until egress had occurred in a majority of schizonts. Image data were analysed using NIS-Elements AR Analysis Software v. 4.51.01. Three replicate experiments recording DG formation to egress provided timing data for 47, 14 and 30 schizonts, respectively. Schizonts were numbered and timing from granular fluorescent patterning formation to egress was measured individually.

### Statistics

Box-and-whisker plot, interquartile interval and average of timing of DG formation to egress times were generated using GraphPad Prism 9.

### Co-expression network analysis

A query of the www.malaria.tools database using RESA (PF3D7_0102200) as input gave an initial co-expression neighbourhood dataset of 59 proteins (Supplementary Table 1). The dataset was further analysed using PlasmoDB.org; proteins lacking a signal sequence or transmembrane (TM) domain or annotated as having a function unlikely to be performed by secreted proteins were removed. The final list of proteins for further investigation comprised 23 proteins (Table 1).

### Immunofluorescence assays

Smears of late-stage segmented 3D7 *P. falciparum* schizonts purified on a Percoll gradient were air dried and stored with desiccant beads at −20°C. Parasites were fixed with acetone for 30 minutes, circled using an immuno-pen and subsequently blocked with 3% BSA in PBS (blocking solution) for a further 30 minutes. Primary antibodies were diluted in blocking solution and applied to the smears which were incubated at room temperature for 1 hour. Anti-HSP40 (PF3D7_0501100) antibodies (a kind gift of Prof. Catherine Braun-Breton, University of Montpellier) were used at a dilution of 1:250; anti-AMA1 and anti-Ron4 antibodies (kind gifts of Mike Blackman, Francis Crick Institute) were used at a dilution of 1:100 and 1:250, respectively, and the anti-RESA monoclonal antibody 28/2 (obtained from the antibody facility at the Walter and Eliza Hall Institute of Medical Research) was used at a dilution of 1:500. The slides were then washed three times with PBS before application of the appropriate fluorophore-linked secondary antibodies and Hoechst 33342 nuclear stain at 15 µg/mL in blocking solution and incubation for a further 45 minutes at room temperature. Following three washes with PBS, the slides were covered with Vectashield antifade mounting medium and sealed with nail polish. Parasites were imaged on a Nikon Eclipse TE fluorescence microscope equipped with a Hamamatsu ORCA-flash 4.0 digital camera C11440. The images were deconvolved using the Richardson-Lucy algorithm with 15 iterations using Nikon NIS-Elements version 5.3 software. The images where then separated, cropped, coloured and overlaid using FIJI software. Images were further cropped and sized using Photoshop. Figures were produced using Illustrator. Pearson’s coefficients were generated using FIJI (ImageJ) Colocalize tool.

## RESULTS

### Generation of a reporter strain for the study of dense granule formation

To study the biosynthesis of DGs and the movement of DG proteins through the secretory system, we fused the gene encoding the well-established DG marker RESA (Culvenor et al., 1991) with the gene encoding mNeonGreen (mNG). This fusion was introduced into 3D7 parasites and integration into the native *RESA* locus was selected for using selection-linked integration (SLI) (Birnbaum et al., 2017b) (Figure 2A). Integration of the RESA-mNG gene fusion into the native RESA locus was verified using PCR and production of the expected mNG fusion protein product was verified by immunoblotting (Figures 2B and 2C). The transgenic parasites were brightly fluorescent in the schizont stage and displayed the expected fluorescence around the periphery of the infected erythrocyte after the parasites had invaded, as seen previously with RESA-GFP fusions (Rug et al., 2004) (Figure 2D).

**Figure 1.**
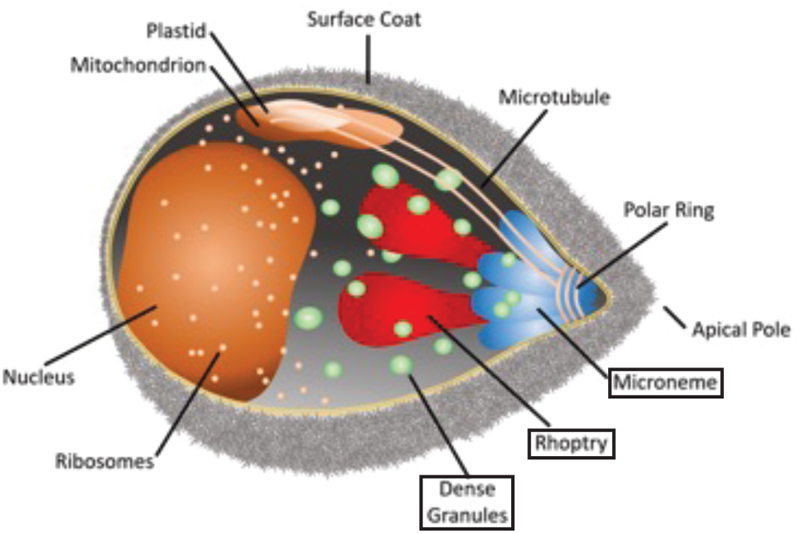
A *Plasmodium falciparum* merozoite. Indicated are the different organelles inside the parasite, with the names of the apical organelles in boxes.

**Figure 2.**
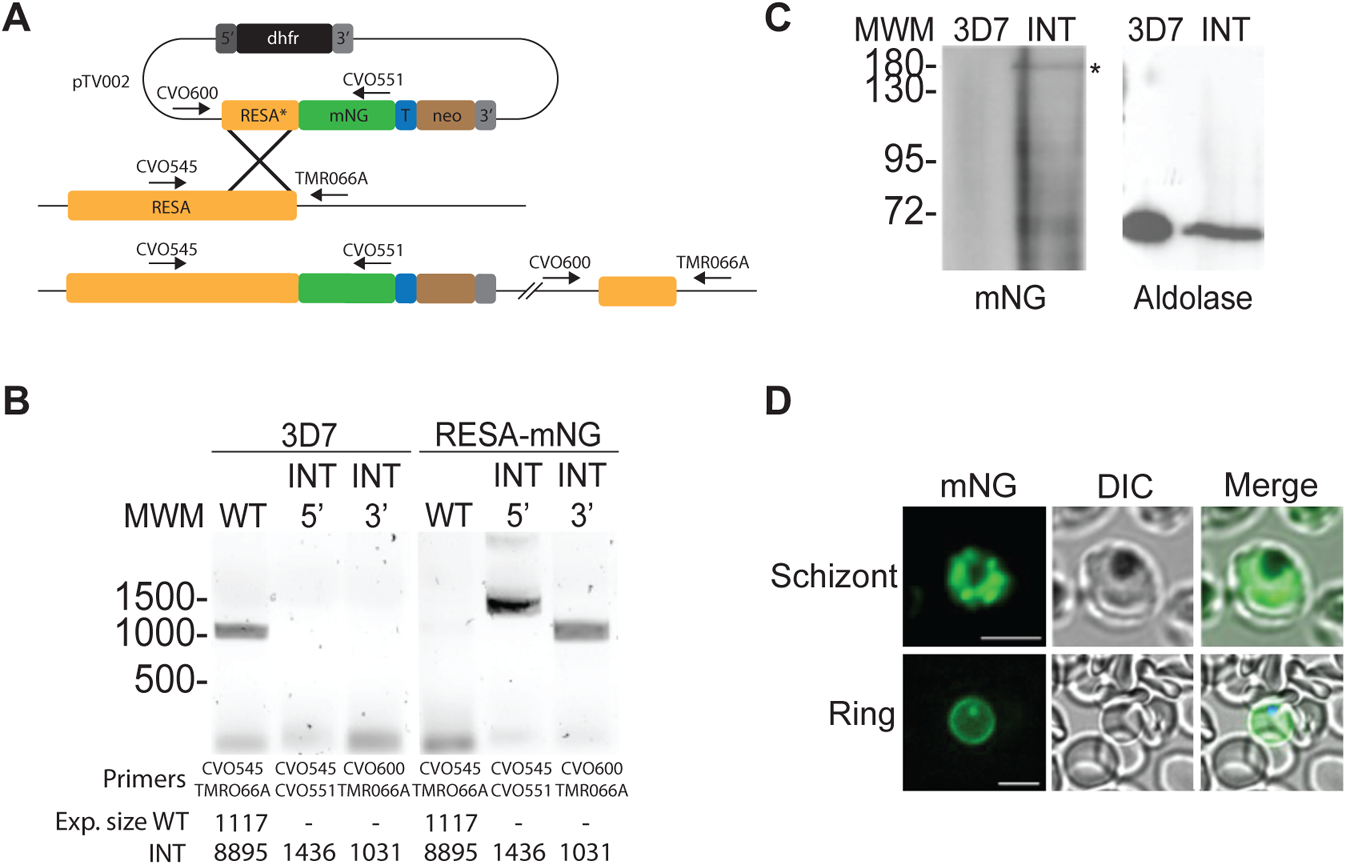
Generation of an mNeonGreen-RESA-expressing parasite. A) Integration strategy using selection-linked integration (SLI). mNG-mNeonGreen; T-T2A peptide; neo-amino 3′-glycosylphosphotransferase (neomycin resistance gene); 5’ and 3’: 5’ and 3’ regulatory regions, respectively. The plasmid pTV002, containing a fusion of the 3’ region of the RESA gene and the gene encoding mNG, was introduced into 3D7 parasites. Transfectants were selected with WR and integrants were subsequently selected with G418. B) Integration PCR of genomic DNA from wildtype (3D7, left) and integrant (INT, right) parasites using the indicated primer pairs. See panel A for the binding sites of the primers. The expected sizes of the PCR products are indicated at the bottom. C) Anti-mNeonGreen immunoblot of 3D7 (right) and RESA-mNG integrant parasite extracts (left). The band of the expected size is indicated with (*). The same extracts were probed with anti-aldolase antibodies as loading control (right). D) Live-cell fluorescence imaging of parasites expressing RESA-mNG. Top-schizont; bottom–ring. Scale bars: 5 µm.

### Timing of dense granule biogenesis

To determine the timing of DG biogenesis we observed the appearance of green fluorescence and the coalescence of the fluorescence into distinct organelles using live video microscopy imaging of the transgenic RESA-mNG-expressing parasites. As RESA expression initiates late in the erythrocytic cycle, as supported by our initial imaging experiments, we started imaging tightly synchronised late-stage schizonts from 46-47 hours post-invasion at five-minute intervals. A five-minute interval was chosen to minimise the toxic effect of the laser exposure whilst providing an interval narrow enough to provide a reliable time frame. In three replicate experiments, schizonts were observed to undergo DG biogenesis, from the appearance of DG fluorescence to egress. The start of DG biosynthesis was set at the time of the appearance of granular foci of fluorescence (Figure 3A). The averaging of the times within the interquartile interval of each replicate revealed that DG formation occurs approximately 37 minutes prior to egress from the iRBC (Figure 3B). Several outliers were observed in which the period from DG formation to egress was far greater or shorter than the majority of the other events. These outliers, which fell outside of the interquartile intervals, were not included in the calculation of average DG formation to egress timing. In the outliers in which the timing of DG formation to egress greatly exceeded the average, the cells were observed to reach a point at which egress would be expected to occur based on appearance by both fluorescence and DIC microscopy, but egress appeared blocked or delayed. In these instances, DG fluorescence grew progressively brighter than in other cells as the RESA-mNG fusion continued to be produced and transported to the DGs. We had observed this phenomenon previously in cells exposed to higher laser intensities and in the absence of the antioxidant resveratrol, in which we found that imaging using laser intensity over 5% and without the antioxidant resveratrol inhibited egress and lead to eventual parasite death.

**Figure 3.**
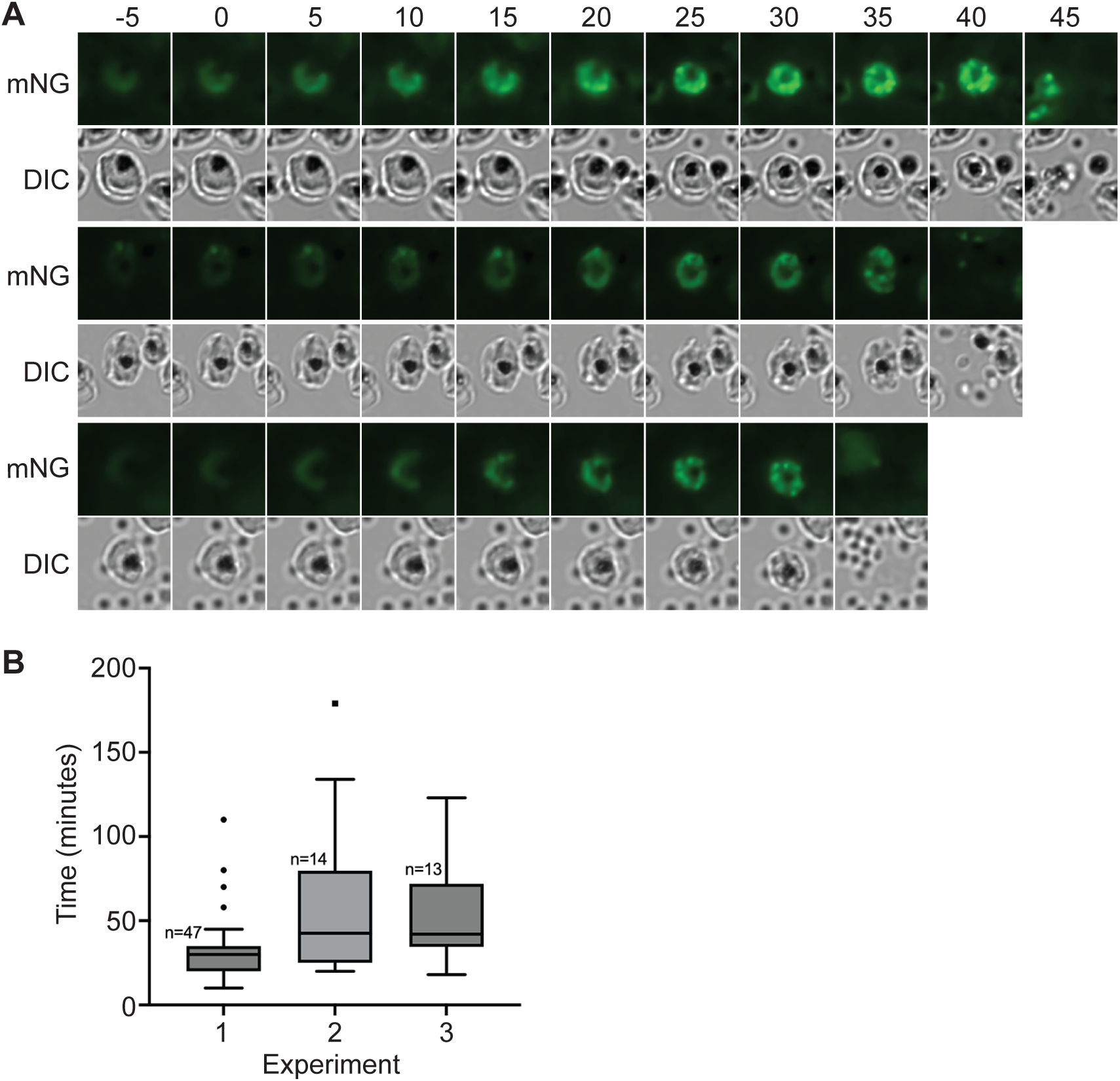
Video microscopy of dense granule formation. A) Highly synchronized RESA-mNG expressing parasites were observed using live video microscopy at intervals of five minutes. Parasites were imaged on three separate occasions; each row is taken from a different experiment. The formation of punctate spots of fluorescence was designated as the start of dense granule formation (time point 0). mNG-mNeonGreen fluorescence; DIC-differential interference contrast. B) Quantitation of dense granule formation for the three different experiments. The boxes represent the interquartile range, where 50% of the data points are found. The horizontal line crossing the box represents the median.The y axis represents the time (in minutes) from the first detection of clustering of RESA-mNG fluorescence in spots to egress of the parasites.

As *P. falciparum* rhoptries are formed predominantly between the 2^nd^ and 4^th^ round of nuclear division and merozoite formation begins at the end of the 4^th^ round of nuclear division (Margos et al., 2004), these experiments indicate that DGs are unusual among the apical organelles in that they are formed well after the final round of nuclear division – on average only 37 minutes prior to egress – a tiny fraction (1.3%) of the entire 48-hour intraerythrocytic life cycle.

### Co-expression network analysis to identify additional dense granule proteins

As our results above revealed that DGs are formed very late in the intraerythrocytic cycle, we inferred that expression network analysis to find proteins with a late-stage expression peak around the time of DG formation could identify potential additional DG proteins for further investigation. We therefore searched for proteins with expression profiles similar to that of RESA using the malaria.tools database (Tan & Mutwil, 2020) (Figure 4A). This provided a co-expression neighbourhood dataset comprising 59 proteins (Supplementary Table 1). A query of Plasmodb.org of the identified proteins provided data on protein features, including TM domains, signal sequences and predicted export signals, mutant phenotype and rodent genetic modifications where available (Amos et al., 2022; David S & Roos, 2001). Of the proteins in the original malaria.tools output, 39 (66%) contained one transmembrane domain or a signal sequence, indicating that they can enter the secretory pathway. The output also contained 6 RESA orthologues, which were included for further analysis. Proteins with an annotated function that indicates that they are not present in DG were omitted.

**Figure 4.**
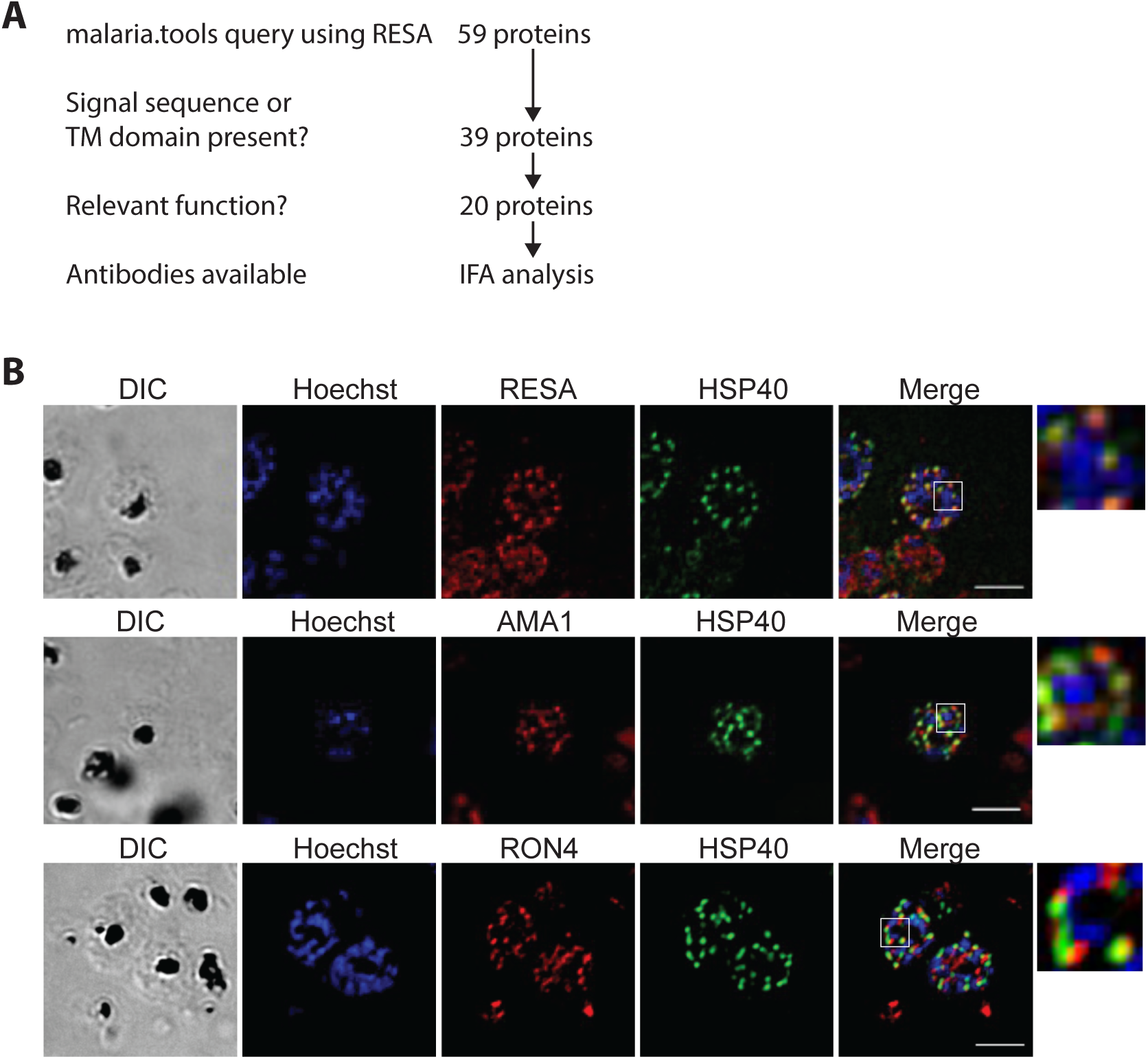
Identification of additional dense granule proteins. A) Outline of malaria.tools query using RESA (PF3D7_0102200). B) Co-staining of *Plasmodium falciparum* parasites using anti-HSP40 and the dense granule marker RESA (top), the microneme marker AMA1 (middle) and the rhoptry neck marker RON4 (bottom). Samples were also stained with Hoechst 33342 to visualize DNA. Panels on far right show an expanded view of the region indicated in the white boxes. Scale bar represents 5 µm.

Proteins were selected for further analysis on the criteria of having a previously described function with potential relevance to erythrocyte remodelling or protein transport, containing a signal sequence or TM domain capable of targeting the protein to the secretory pathway, and having antibodies available for IFA analysis. Selection based on these criteria gave a list of 20 potential DG proteins (Table 2). Of the proteins that did not contain a TM domain or signal sequence, three were included in the final set based on their predicted function (v-SNARE, gametogenesis implicated protein (GIG), and DYN3/DrpC). As v-SNAREs mediate fusion of vesicles and granules to target membranes (Chen & Scheller, 2001; Dhara et al., 2016; Wang et al., 2017), this v-SNARE was included to investigate a potential role in DG protein transport or fusion of DGs to the parasite membrane. GIG is implicated in gametocyte production and was included as it may point to gametocyte-specific alteration of the host-cell (Gardiner et al., 2005). The dynamin-like protein DrpC was included as tgDrpC is involved in vesicular transport and mitochondrial fission (Heredero-Bermejo et al., 2019; Melatti et al., 2019) (Breinich et al., 2009), and tgDrpB is required for formation of secretory organelles in *T. gondii* (Spielmann et al., 2020). The role of DrpC in endocytosis has not been investigated. The final list of putative DG proteins comprised 23 proteins (Table 2). The previously identified DG proteins PTEX88 and LSA3 are present in the malaria.tools output, as well as several RESA N-terminal sequences, supporting the use of this technique as a tool for identifying DG proteins. However, as certain DG proteins, including EXP1 and EXP2, do not have this distinctive late-stage expression peak they were not present in the output and it is likely that other DG proteins were not identified in this analysis for the same reason. Exported proteins are greatly overrepresented in the final list, and proteins for which mutant phenotypes have been described, for example PTP1 and 5, and members of the FIKK family, predominantly have roles in host cell remodelling and cytoprotection (Davies et al., 2020; Maier et al., 2008; Rug et al., 2014). These characteristics are in keeping with the model that DGs are important mediators of host cell remodelling.

**Table 2.**
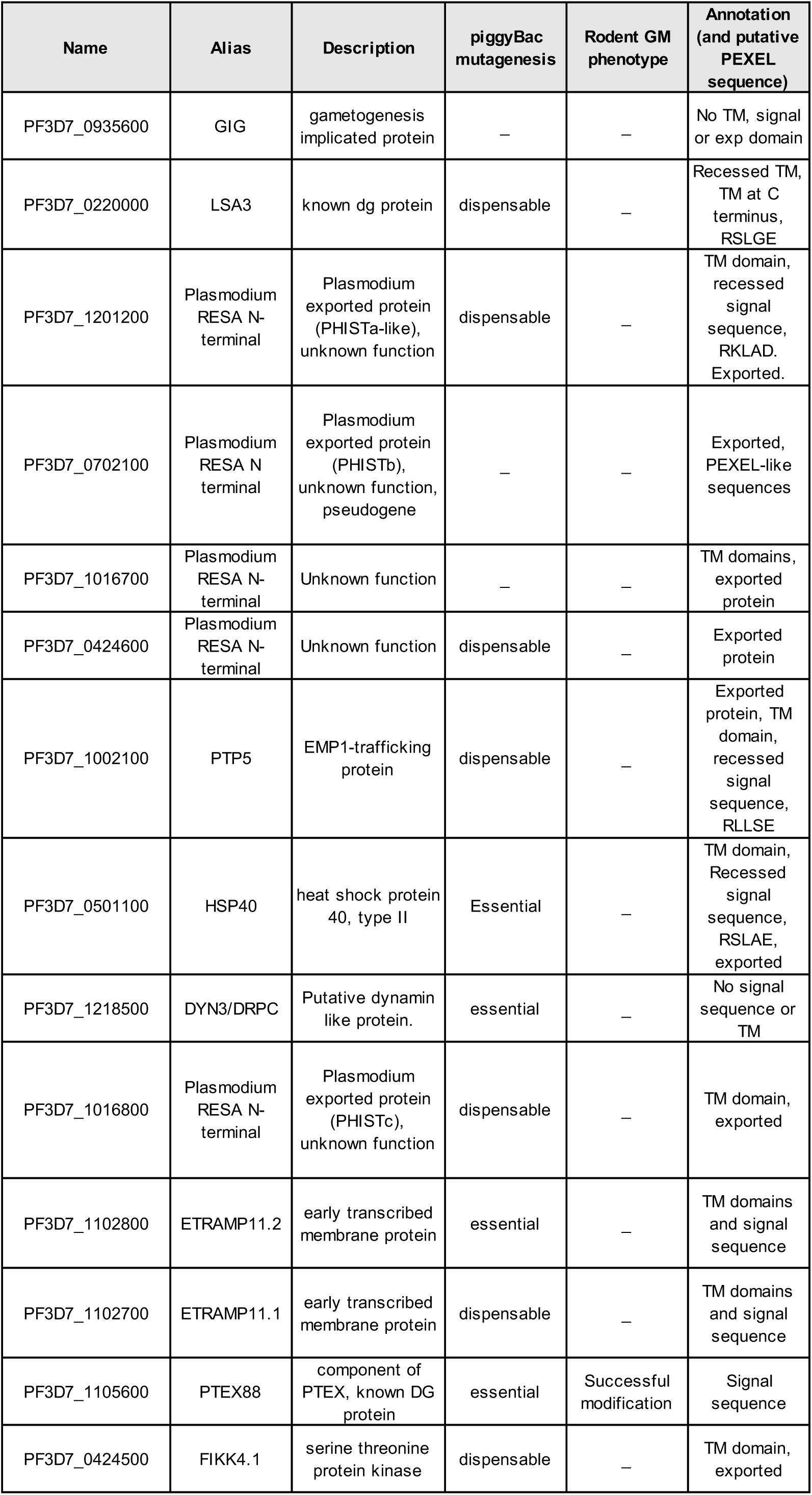
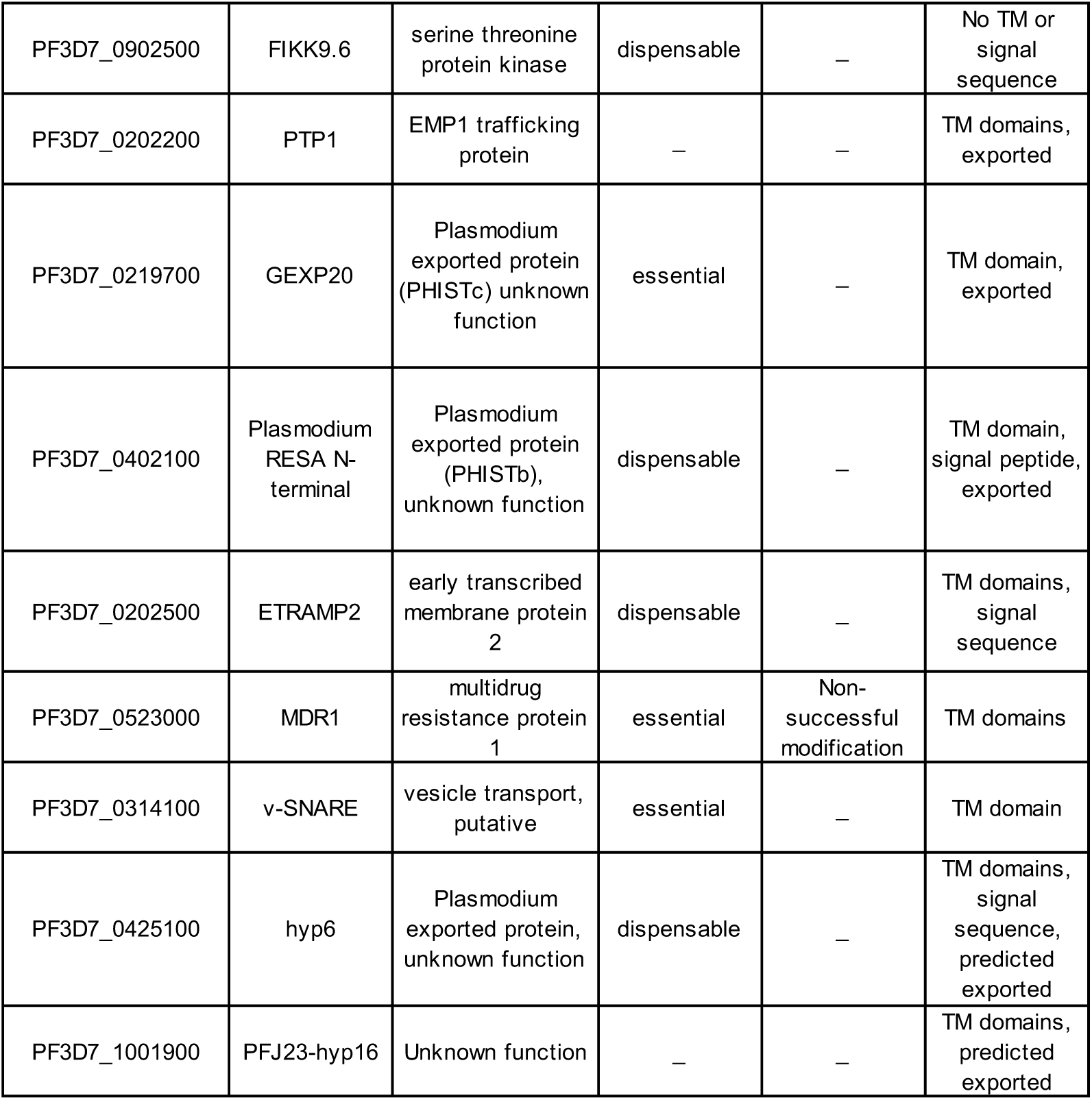
Putative *Plasmodium falciparum* dense granule proteins identified in malaria.tools using RESA as query

### Verification of localisation of putative dense granule proteins

To verify whether protein identified in the bioinformatics screen are transported to DGs, we determined the localization of one of the potential DG proteins, HSP40, by IFA. Co-staining of late-stage schizonts with microneme, rhoptry and DG markers (AMA1, RON4 and RESA, respectively) revealed strong overlap of the RESA and HSP40 signal in all late-stage segmented schizonts, indicating co-localisation of the two proteins (Figure 4B). Indeed, no examples of HSP40 staining distinct from RESA staining was detected in late-stage segmented schizonts. Staining using antibodies against the rhoptry neck marker Ron4 and the microneme marker AMA1 exhibit greatly lower levels of colocalization with anti-HSP40 antibodies. In these samples HSP40 staining was detected distinctly adjacent to AMA1 and RON4 staining. Strong overlap with DG marker staining compared to the distinct staining patterns seen with microneme and rhoptry markers indicate that HSP40 localises to DGs and not to the micronemes and rhoptries. This was further supported by the Pearson’s correlation coefficient (PCC) for the antibody pairs HSP40-RESA (0.810), HSP40-AMA1 (0.370) and HSP40-RON4 (0.205) (Supplementary Figure 1). This result identifies an additional DG protein and verifies the use of co-expression network analysis as a tool for DG protein identification.

## DISCUSSION

Here we have shown that DGs in *Plasmodium falciparum* are formed extremely late in the erythrocytic life cycle. In our experiments DGs were detected on average 37 minutes prior to egress. However, as egress timing appears to be delayed by the oxidative stress induced by laser exposure and mNG fluorescence, this estimation of the start of DG biogenesis is likely to be an underestimation. Hence, DG biogenesis is a very rapid process in which the organelles are synthesized very quickly.

As DGs are formed very late in the intraerythrocytic cycle and several known DG proteins, including RESA, consequently have a distinctive late-stage expression peak, we inferred that expression network analysis to find other proteins with a similarly distinctive late expression peak could identify potential DG proteins for further investigation. Through a search of the malaria.tools co-expression network platform using RESA as a query we identified genes with expression profiles similar to that of RESA, including heat shock protein HSP40. We did indeed find that HSP40 co-localizes with RESA, indicating that HSP40 is a DG protein (Liu et al., 2020). This supports the use of co-expression network analysis as a method for DG protein prediction. As HSP40 is part of the J-dots, which are small parasite-derived structures found in the cytosol of infected erythrocytes that may be involved in the transport of parasite proteins through the host cell cytosol (Külzer et al., 2010; Petersen et al., 2016), it indicates that at least some the components of these structures are present in the DGs and exported to the host cell almost immediately after the parasite enters the host cell. Although the function of these structures remains unclear, it has been postulated that they may have a function in the transport of parasite proteins through the erythrocyte cytosol (Behl et al., 2019). Hence, by releasing the component protein of J-dots in DGs, the parasite may enhance its ability to target these proteins to their proper intraerythrocytic location almost immediately after invasion. This finding therefore may also further support the hypothesis that DGs likely contain many more exported proteins that are yet to be identified. This aligns with the timing of DG discharge immediately following invasion and the hypothesised role of DGs in erythrocyte remodelling. Interestingly, one protein identified in the malaria.tools investigation, PTP1, is necessary for correct formation of the Maurer’s clefts and linking the Maurer’s clefts to the cytoskeleton (Rug et al., 2014). This may well indicate that Maurer’s clefts are formed very soon after invasion, using proteins that are exported immediately after DGs have been released.

Our bio-informatics approach identified the known DG proteins LSA3 and PTEX88 (Bullen et al., 2012; Morita et al., 2017). However, several other DG proteins, including EXP1 and EXP2, were not included in the malaria.tools output, likely as they do not share the distinctive late-stage expression peak of RESA and instead are also expressed at other times throughout the life cycle. It is likely that other unidentified DG proteins will also be missing from the output owing to having expression profiles dissimilar to that of RESA. Combined, the listing of known and recently identified DG proteins, and the localisation of the one protein that has been tested to date to the DGs suggests that co-expression network analysis using malaria.tools works as a predictive method of identifying candidate proteins for further analysis. Further, exported proteins are over-represented within the dataset. As host cell effector proteins must be exported, the high number of exported proteins within the dataset accords with the understanding of the role of DGs as a secretory compartment containing proteins with roles in host cell modification that are released immediately after erythrocyte invasion. A recently described database of DG protein in *Plasmodium* proteins also predicts that several PHIST proteins are present in DGs (Hu et al., 2022).

Three proteins with a transcriptional profile similar to that of RESA are of potential interest based on their putative function, despite lacking a signal sequence or TM domain: PF3D7_0314100, annotated as a v-SNARE GiG and the dynamin-like protein DrpC. These proteins may serve important functions in the fusion of the DGs to the parasite plasma membrane, gametocyte-specific functions immediately after invasion and vesicular trafficking to the DGs, respectively (Bai et al., 2018; Chen & Scheller, 2001; Dhara et al., 2016; Gardiner et al., 2005; Heredero-Bermejo et al., 2019; Melatti et al., 2019; Wang et al., 2017).

This is the first description of the timing of DG biogenesis in *Plasmodium* parasites. This finding reveals that DGs are only present within the parasite for a very short portion (1.3%) of the 48-hour asexual cycle. We also find that the exported protein HSP40 is present in DGs. As HSP40 interacts with HSP70 to form a chaperone complex within the infected host cell, this finding further supports the hypothesis that DGs have an important role in host cell modification, in this instance through the chaperoning of parasite effector proteins (Liu et al., 2020). These new insights shed further insight into how the parasite prepares for modification of the host erythrocyte starting very late in the schizont phase.

## ACKNOWLEDGEMENTS

We are grateful to Dr James Thomas (LSHTM) for providing the JT02-01-31 plasmid containing the SLI fragment and Dr Gerhard Wunderlich (Institute of Biomedical Sciences, University of Sao Paulo) for providing plasmid pRESA-HA. We thank Prof. Catherine Braun-Breton (University of Montpellier) for providing the anti-HSP40 antibody, and Mike Blackman (Francis Crick Institute) for the anti-AMA1 and anti-RON4 antibodies. The authors acknowledge the facilities and the scientific and technical assistance of the LSHTM Wolfson Cell Biology Facility, with specific thanks to Liz McCarthy. This work was supported by an MRC-LID studentship to TV and a Medical Research Council Career Development Award (MR/R008485/1) to CvO.

---

Supplementary Figure 1. **Quantitation of colocalization**. Pearson’s correlation coefficient (PCC) of the colocalization of the anti-HSP40 staining and the staining using the indicated antibodies that recognize apical organelle markers. For each combination, the PCC was determined using ten clearly labeled merozoites in different schizonts.

