## Supplementary figures and images for "Timing of dense granule biogenesis in asexual malaria parasites"

### Supplementary Figure 1

Supplementary Figure 1

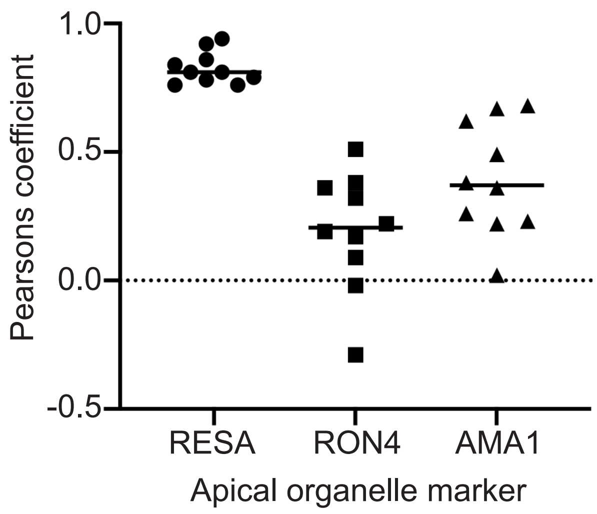
