## Supplementary Table 1 for "Timing of dense granule biogenesis in asexual malaria parasites"

| | |
|---|---|
| PF3D7_1115800 | |
| PF3D7_0701900 | |
| PF3D7_0935600 | GIG |
| PF3D7_0220000 | LSA3 |
| PF3D7_1201200 | Plasmodium RESA N-terminal. |
| PF3D7_0702100 | Plasmodium RESA N-terminal. |
| PF3D7_1002100 | PTP5 |
| PF3D7_0501100 | HSP40 |
| PF3D7_1218500 | DRPC |
| PF3D7_1016800 | Plasmodium RESA N-terminal. |
| PF3D7_0301800 | |
| PF3D7_0605900 | P GNS1/SUR4 |
| PF3D7_0102700 | MaTrA |
| PF3D7_0730800 | |
| PF3D7_0814500 | |
| PF3D7_1114200 | Rab-GTPase-TBC domain. |
| PF3D7_0525100 | SSF56801: AMP-binding enzyme. |
| PF3D7_1102800 | ETRAMP11.2 |
| PF3D7_1102700 | ETRAMP11.1 |
| PF3D7_0108500 | ELM2 domain. |
| PF3D7_1010300 | SDH4 0 |
| PF3D7_0606000 | |
| PF3D7_0702000 | |
| PF3D7_0830500 | TryThrA |
| PF3D7_1252600 | SSF53474:"Serine aminopeptidase, S33". 0.6779441589522898. esterase, putative |
| PF3D7_1105600 | PTEX88 |
| PF3D7_1343700 | SSF117281: Kelch13 |
| PF3D7_1237900 | |
| PF3D7_0424500 | FIKK4.1 |
| PF3D7_1200700 | SSF56801: No description available. Pfam domain(s): PF00501: AMP-binding enzyme. ACS7 0.6710509489350647. acyl-CoA synthetase. |
| PF3D7_0902500 | SSF56112: Protein kinase-like domain superfamily. Pfam domain(s): PF00069: Protein kinase domain. FIKK9.6 0.6674477893666778 |
| PF3D7_1149600 | SSF46565: Chaperone J-domain superfamily. Pfam domain(s): PF00226: DnaJ domain, PF14308: X-domain of DnaJ-containing. 0.6618812038153901 |
| PF3D7_1016700 | No superfamily available. Pfam domain(s): PF09687: Plasmodium RESA N-terminal. 0.6589725695663241 |
| PF3D7_0423900 | No superfamily available. Pfam domain(s): No Pfam domain available. 0.6507065747644319 |
| PF3D7_0202200 | No superfamily available. Pfam domain(s): No Pfam domain available. PTP1 0.6467386611328264 |
| PF3D7_0805200 | No superfamily available. Pfam domain(s): No Pfam domain available. GAMER 0.6462594786755821 |
| PF3D7_1416500 | SSF51735: NAD(P)-binding domain superfamily, SSF53223: No description available. Pfam domain(s): PF00208: Glutamate/Leucine/Phenylalanine/Valine dehydrogenase, PF02812: "Glu/Leu/Phe/Val dehydrogenase, dimerisation domain". GDH1 0.6409844721556132 |
| PF3D7_1001100 | SSF47027: Acyl-CoA binding protein superfamily. Pfam domain(s): PF00887: Acyl CoA binding protein, PF09687: Plasmodium RESA N-terminal. ACBP1 0.638974250206304 |
| PF3D7_0731100 | Plasmodium RESA N-terminal. GEXP20 |
| PF3D7_0402100 | No superfamily available. Pfam domain(s): PF09687: Plasmodium RESA N-terminal. 0.629191155827773 66 PF3D7_0402200 No superfamily available. Pfam domain(s): No Pfam domain available. SURF4.1 0.6247482190782291 |
| PF3D7_1479000 | SSF56801: No description available. Pfam domain(s): PF00501: AMP-binding enzyme. ACS1a 0.6189628527678525 |
| PF3D7_1433500 | SSF56719: "DNA topoisomerase, type IIA-like domain superfamily", SSF55874: Histidine kinase/HSP90-like ATPase superfamily, SSF54211: Ribosomal protein S5 domain 2-type fold. Pfam domain(s): PF00521: "DNA gyrase/topoisomerase IV, subunit A", PF01751: Toprim domain, PF16898: C-terminal associated domain of TOPRIM, PF02518: "Histidine kinase-, DNA gyrase B-, and HSP90-like ATPase", PF00204: DNA gyrase B. TOP2 0.6176353572557606 |
| PF3D7_0202500 | No superfamily available. Pfam domain(s): PF09716: Malarial early transcribed membrane protein (ETRAMP). ETRAMP2 0.6167221285901517 |
| PF3D7_0219900 | No superfamily available. Pfam domain(s): No Pfam domain available. 0.6130706441449282 |
| PF3D7_1404800 | No superfamily available. Pfam domain(s): No Pfam domain available. 0.6114481970203258 |
| PF3D7_1478800 | No superfamily available. Pfam domain(s): PF09715: Plasmodium protein of unknown function (Plasmod dom 1). 0.6084027133613183 |
| PF3D7_0523000 | SSF90123: "ABC transporter type 1, transmembrane domain superfamily", SSF52540: P-loop containing nucleoside triphosphate hydrolase. Pfam domain(s): PF00005: ABC transporter,PF00664: ABC transporter transmembrane region. MDR1 0.5995779354021784 |
| PF3D7_1404900 | No superfamily available. Pfam domain(s): No Pfam domain available. 0.5866462420187923 |
| PF3D7_1026600 | No superfamily available. Pfam domain(s): No Pfam domain available. 0.5768375433341388 |
| PF3D7_0314100 | SSF58038: No description available. Pfam domain(s): PF05008: Vesicle transport v-SNARE protein N-terminus. 0.5745987799771859 |
| PF3D7_1001900 | No superfamily available. Pfam domain(s): No Pfam domain available. PfJ23 0.5709363884342216 |
| PF3D7_1352900 | No superfamily available. Pfam domain(s): No Pfam domain available. 0.558809219660643 |
| PF3D7_0702200 | SSF53474: Alpha/Beta hydrolase fold. Pfam domain(s): PF12146: "Serine aminopeptidase, S33". 0.5449531215890748 |
| PF3D7_1226300 | SSF56784: HAD-like superfamily. Pfam domain(s): PF08282: haloacid dehalogenase-like hydrolase. HAD2 0.5340504460320102 |
| PF3D7_1203700 | SSF143113: NAP-like superfamily. Pfam domain(s): PF00956: Nucleosome assembly protein (NAP). NAPL 0.5319418943357009 |
| PF3D7_1477800 | SSF47027: Acyl-CoA binding protein superfamily. Pfam domain(s): PF00887: Acyl CoA binding protein. ACBP 0.5275781495997893 |
| PF3D7_0425100 | No superfamily available. Pfam domain(s): PF09715: Plasmodium protein of unknown function (Plasmod dom 1). 0.5045612465857853 93. hyp6 |
