## Supplementary Table 2 for "Timing of dense granule biogenesis in asexual malaria parasites"

**Supplementary Table 2** Primers used in this study

| Name | Sequence | Restriction site |
|---|---|---|
| CVO545 | GTGAACAAATGAATTCAATAACATACAATTTCG | none |
| CV0550 | GGAC**CTGCAG**AAAAAGTTAGTAAAGGAGAAGAAGATAATATGGCAAG | PstI |
| CVO551 | GGAC**ACGCGT**TGCTTTATACAATTCATCCATTCCCATAACATCTGTAAATG | MluI |
| CVO576 | GGAC**ACGCGT**GAAGGAAGAGGAAGTTTATTAACATGTGGAG | MluI |
| CVO577 | GGAC**ACGCGT**TTAGAAGAACTCGTCAAGAAGGCG | MluI |
| CVO600 | CAAAATGGTTAACAAAGAAGAAGCTCAGAG | none |
| TMR066A | CACATTTCTTTTTCATATATCTTGTTTTAAATATTTTATTTTCATAAAGAG | none |
